# A tumor-suppressive role of the PRC1 Polycomb epigenetic complex in the maintenance of adult *Drosophila* intestinal stem cell identity

**DOI:** 10.1101/2025.03.18.643963

**Authors:** Aurélia Joly, Ana-Maria Popmihaylova, Corinne Rancurel, Julie Soltys, Rihab Loudhaief, Armel Gallet, Edan Foley, Anne-Marie Martinez, Raphaël Rousset

**Affiliations:** Université Côte d’Azur, INRAE, ISA, France; Université Côte d’Azur, CNRS, INRAE, EMR PHYBAC, France; Department of Medical Microbiology and Immunology, University of Alberta, Edmonton AB, Canada; Institute of Human Genetics, Univ. Montpellier, CNRS, Montpellier, France

**Keywords:** Epigenetics, Polycomb, tumor suppression, intestinal stem cells, *Drosophila*

## Abstract

Chromatin modulators, like Polycomb group proteins, are key epigenetic regulators of gene expression and are frequently mutated in cancers. In adult stem cells, epigenetic regulation maintains their identity and controls their differentiation during homeostasis or aging, but its direct role in tumorigenesis remains unclear. Here we developed a novel tumor model in *Drosophila* by exploring the function of Polycomb Repressive Complex 1 (PRC1) in adult intestinal stem cells (ISCs). Disrupting core PRC1 components in ISCs induces the formation of small cell clusters devoid of intestinal markers, a novel phenotype linked to premature mortality under stress. These clusters exhibit neoplastic characteristics such as overproliferation and continuous growth in serial transplantations, leading to their designation as tumor-initiating intestinal cells (TIICs). While JAK/STAT signaling contributes to TIIC growth, the NF-κB-related Toll/Imd immune pathways restrict their expansion independently of cell death. Altogether, our results highlight PRC1 as an epigenetic tumor suppressor in adult stem cells.

**AUTHOR SUMMARY:** Our research explores how stem cells in the adult intestine stay healthy and avoid becoming cancerous. We focused on a group of proteins called Polycomb Repressive Complex 1 (PRC1), which help regulate which genes are turned off in a cell. While these proteins are known to play important roles during development and cancer prevention, their function in the adult intestine has been less clear. Using the fruit fly *Drosophila*, we discovered that when PRC1 function is lost in intestinal stem cells, abnormal clusters of cells begin to form. These clusters grow uncontrollably, lose their normal identity, and can keep growing when transplanted—traits that are typical of cancer. Interestingly, we also found that parts of the immune system, specifically NF-κB-related pathways, can act inside these tumor cells to limit their growth—revealing a protective role that hasn’t been seen before in the adult gut. This work provides new insights into how epigenetic regulation and immune signaling work together to keep stem cells from turning cancerous. It opens up new possibilities for understanding how cancers begin and how the body may naturally resist them, even at the level of the tissue.

## INTRODUCTION

Adult stem cells balance self-renewal and differentiation by replacing damaged specialized cells according to tissue needs [1]. Equally important, proliferation is regulated to avoid excess cell production leading to hyperplasia and tumorigenesis. Transcription programs and their dynamic changes must be finely controlled to adjust stem cell responses, and how these changes are regulated remains a fundamental question. Chromatin organization into highly dynamic domains modulates transcription programs in a cell type-dependent manner [2, 3]. During differentiation, stem cells reshape this organization to enhance cell type-specific transcription program and achieve specialized cellular functions [4]. Whether and how these dynamic changes regulate stemness and differentiation of adult intestinal stem cells is poorly explored. Since dysregulation of adult stem cells is associated with a variety of diseases like cancer, it appears essential to study the mechanisms that control adult stem cell behavior.

Polycomb group (PcG) proteins are major regulators of transcription programs that shape three-dimensional genome organization. Initially characterized for their role in conserved developmental processes, they have also been implicated in various mature processes, notably in adult stem cell regulation [5, 6]. Their crucial importance and efficiency led to a high conservation of these proteins and their mechanisms from plants to mammals. PcG proteins combine into multi-protein complexes to bind and shape chromatin and thus transcription programs through the deposition of post-transcriptional histone modifications [7]. In the fruit fly *Drosophila melanogaster*, the core canonical PRC1 includes Sex combs extra/Ring (Sce/dRing), Posterior sex combs (Psc), Polyhomeotic (Ph), and Polycomb (Pc), a chromodomain-containing protein that recognizes the trimethylation of histone H3 at lysine27 (H3K27me3) mark deposited by the Polycomb Repressive Complex 2 (PRC2) [6]. The core PRC1 E3 ligase catalytic subunit Sce/dRing requires interaction with Psc, and its paralog Suppressor of zeste 2 (Su(z)2), to form an active PRC1 core and to ubiquitinate lysine 118 (lysine 119 in mammals) of histone 2A (H2AK118ub). The mammalian orthologues of Sce and Psc are RING1A/B and PCGF1-6, respectively [8]. Misregulation of PcG protein levels has been linked to various human cancers [9, 10]. Bmi-1 (PCGF4) was initially identified as an oncogene that collaborates with c-Myc in murine lymphoma [11]. In contrast, another Psc homolog, Mel-18 (PCGF2), has been characterized as a tumor suppressor, notably through transcriptional repression of *Bmi-1* [12, 13]. Several studies in vertebrates have examined the role of PcGs in adult stem cells of different tissues, including the epidermis, hematopoietic cells, bone marrow, teeth and intestine [5]. So far, none of these studies have demonstrated that loss of PcGs results in tumor formation. In *Drosophila*, however, the PRC1 subunits Psc and Su(z)2 have been reported to restrain stem cell proliferation in the ovary and testis [14, 15], but their role in other adult stem cells remains unknown.

Intestinal models have played a prominent role in tumor biology, significantly enhancing our understanding of cellular interactions within the tumor microenvironment. The gut epithelium is challenged daily by food intake, microbes, and chemicals, making it one of the most regenerative organs. Most differentiated cells live only a few days in vertebrates, although turnover of the *Drosophila melanogaster* midgut can take three weeks [16–18]. During homeostasis, adult intestinal stem cells (ISCs) ensure the complete replenishment of the diverse specialized intestinal cells required for digestive, absorptive, immune, and endocrine functions [19]. In the *Drosophila* midgut, ISCs divide mostly asymmetrically to self-renew and give rise to progenitor cells: enteroblasts (EBs) and enteroendocrine precursors (EEps) [1]. EBs differentiate into large polyploid absorptive enterocytes (ECs), whereas EEps generate the secretory lineage with enteroendocrine cells (EEs) [20–25]. ISC renewal and differentiation adjust for growth, homeostasis, or regeneration through the coordination of numerous signaling pathways (EGFR, Notch, JAK/STAT, JNK, Wg/Wnt, Dpp, Hippo), as well as integrin and cadherin-based cell-cell adhesions [26, 27]. The proliferation rate and proper differentiation programs must be accurately coordinated, as ISC deregulation is at the origin of intestinal dysfunctions and cancers [28–31]. The dynamics of the transcription program in ISCs are crucial to maintain homeostasis in high-turnover tissues. A previous study in adult *Drosophila* showed that the Pc subunit of PRC1 maintains appropriate specification of ISCs into the EE or EC lineage and that increased Pc activity during aging favors EE cell specification [32]. In the present study, we investigated the role of the other subunits of PRC1 to better understand the epigenetic mechanisms controlling *Drosophila* ISCs.

## RESULTS

### Loss of core PRC1 induces tumor-like clusters in adult ISC

To study PRC1 loss of function in the gut, we first used the chromosomal deficiency XL26 encompassing the *Psc* gene and its redundant paralog *Su(z)2* [14, 33]. We generated fluorescently labelled *Psc/Su(z)2* homozygous mutant clones for 14 days using the FLP-FRT-based MARCM (Mosaic Analysis with a Repressible Cell Marker) system [34]. With this technique, *Psc/Su(z)2* clones are derived from a single mitotic *escargot* (*esg)*-positive progenitor and labelled with GFP. The differentiated cells, ECs and EEs, were distinguished as Prospero-negative large cells (only polyploid cells with large nuclei) and Prospero-positive small-nucleated cells, respectively. In control, neutral clones, we observed both EC and EE lineages derived from the mitotic stem cells, as expected (Fig. 1A-C). In contrast, the *Psc/Su(z)2* loss-of-function clones consist only of Prospero-negative cells with small nuclei, indicating that they acquire neither absorptive (large nuclei) nor secretory (Pros-positive) identity (Fig. 1B, C). These results show that *Psc/Su(z)2*-deficient cells (*esg*-positive) fail to differentiate properly in either lineage.

**Figure 1.**
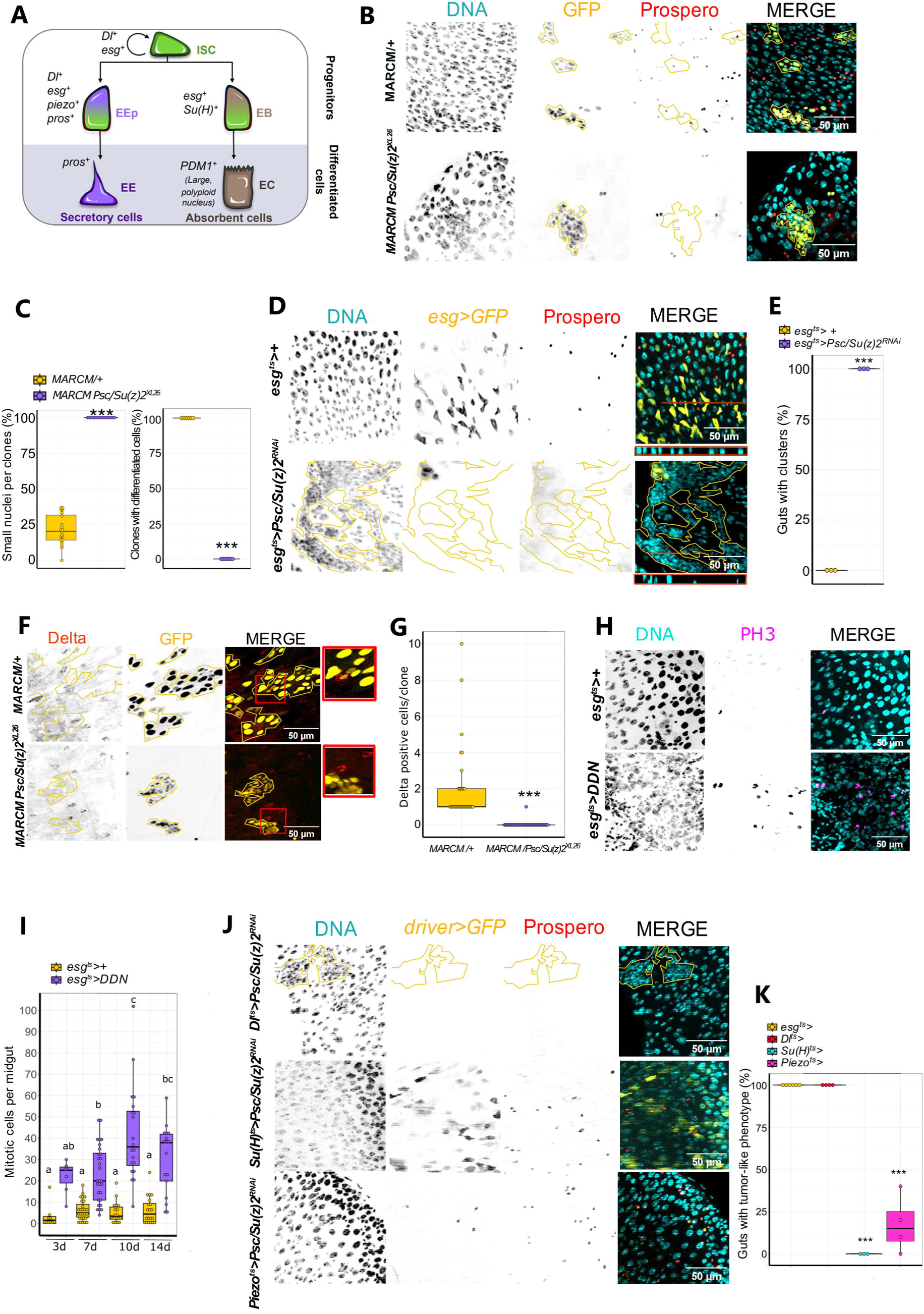
Loss of *Psc/Su(z)2* induces tumor-like clusters in adult ISCs. **A**. Lineage of *Drosophila* intestinal epithelial cells and their specific markers. **B**. Control and *Psc/Su(z)2^XL26^* mutant MARCM clones expressing GFP (yellow). The differentiated EE were marked with the anti-Prospero antibody (red). Clones and clusters are delimited by yellow lines. **C**. Plot on the left shows the percentage (%) of small nuclei in control and mutant clones. *Chi²-test p-value < 2.2e-16. n= 20, one dot represents one measure.* Plot on the right shows the % of the number of clones containing differentiated cells (large nucleus EC and/or Prospero-positive EE). *Chi²-test p-value < 2.2e-16. Septuplicate for a total of n= 203 (control) and 151 (mutant) clones. One dot represents the % in one replicate.* **D**. *esg*-positive cells expressing GFP (yellow) alone (control; upper panels) or together with *Psc/Su(z)2^RNAi^* (bottom panels). Z-projections of the region traced by the brown lines appear below the merged images. **E**. Quantification of guts with tumor phenotype in %. *Anova-Tukey multiple comparisons. One dot represents one replicate with n= 8-12 guts each*. **F**. Control and *Psc*/*Su(z)2^XL26^*mutant MARCM clones expressing GFP (yellow) and stained with the anti-Delta antibody (red). Close-up images of selected clones are also shown (red squares). **G**. Quantification of Delta-positive cells per clone. *Chi²-test p-value < 2.2e-16; n= 37 and 45.* **H**. *esg*-positive cells expressing GFP (yellow) alone (control; upper panels) or with the double dominant negative (DDN) forms of both Psc and Su(z)2 proteins (lower panels). Mitotic cells are stained with the anti-PH3 antibody (purple). **I**. Quantification of PH3 counts per gut from 3d to 14d after induction of expression. *Kruskall-Wallis and Wicolxon test p-values at 3, 7, 10 and 14 days are 0.00028; 5.7e-07; 5.2e-10; 2.2e-08, respectively, for comparison between control and cluster conditions. Different letters indicate that conditions belong to statistically different groups; n= 67, 52, 27, 29, 28, 31, 30,* 25. **J**. Expression of *Psc/Su(z)2^RNAi^*in *ISCs (Dl-GAL4^ts^,UAS-GFP), EBs (Su(H)-GAL4^ts^,UAS-GFP)* and *EEps (Piezo-GAL4^ts^,UAS-GFP)*. **K**. Quantification of guts with tumor phenotype in %. *Anova-Tukey multiple comparisons. One dot represents one replicate of n= 8-12 guts each. Dl-esg = n.s (1.0); Su(H)-Piezo = n.s. (0.0643278); Su(H)-Dl and Su(H)-esg = p <0.00000001; Piezo-Dl and Piezo-esg = p <0.00000001.* DNA (turquoise): stained with DAPI. Scale bar, 50 μm.

To confirm this phenotype, we induced loss of function of both *Psc* and *Su(z)2* either by RNAi (two different lines of *Psc/Su(z)2^RNAi^*) [35] or by co-expression of dominant negative forms of both proteins (Psc/Su(z)2^DDN^,“double dominant negative”) [14] in progenitor cells using the *esg-GAL4^ts^,UAS-GFP* driver. In all cases, we observed epithelial disorganization, visualized by clusters of small nuclei that were Prospero negative and located mainly in the posterior midgut (Fig. 1D, E; Fig. S1A, top three rows). Interestingly, large clusters no longer showed the GFP reporter, suggesting a progressive loss of *esg-GAL4* driver activity. To better follow the cells in which we induced *Psc/Su(z)2* loss of function, we then used an alternative FLP-FRT system: the *esgF/O* line. This line allows cells to permanently express the transgenes (e.g. GFP and RNAi constructs) under the ubiquitous *actin* promoter once recombination is induced by the *esg* promoter. When using this tool with either *Psc/Su(z)2^RNAi^* or *Psc/Su(z)2^DDN^*, we observed large, GFP-positive, Prospero-negative clones with small nuclei that originated from progenitor cells (Fig. S1A, bottom two rows). In agreement with the previous observations, this establishes that *Psc/Su(z)2* loss of function in intestinal progenitors generates clusters of small undifferentiated cells.

The disappearance of GFP reporter gene expression with the *esg-GAL4* driver observed above (Fig. 1D; Fig. S1A, top three rows) suggests that the cell clusters lose expression of the *esg* gene and thus stem cell identity. We tested this hypothesis by analyzing the presence of the ISC specific marker Delta (Dl) in *Psc/Su(z)2* loss-of-function (MARCM or RNAi) cell clusters. As expected, control MARCM clones contain ISCs identified by anti-Dl staining. However, the mutant MARCM clones *Psc/Su(z)2^XL26^* are not composed of any ISCs, as indicated by the absence of Dl expression within the clones (Fig.1 F, G). In accordance with this result, we observed loss of *Dl*-positive cells with *Psc/Su(z)2^RNAi^*or *Psc/Su(z)2^DDN^* (Fig. S1B, top three rows). We also confirmed this result using the *Dl.LacZ* reporter gene (Fig. S1B, fourth row). Occasionally, we observed rare MARCM clusters of two or three cells expressing *Dl*, likely reflecting emerging clones that have not yet lost its expression (Fig. 1G). These results support the notion that *Dl* expression is rapidly lost during cluster formation. We then assessed the expression of *Su(H)*, which is specifically expressed in enteroblasts [36]. The results showed no expression of the *Su(H)-lacZ* reporter in *Psc/Su(z)2^XL26^*MARCM clones, indicating that cell clusters are not composed of enteroblasts (Fig. S1B, fifth row). We also quantified the mitotic activity of the mutant clusters compared to control progenitors using PH3 immunostaining. Control progenitors have a basal level of mitotic activity, allowing a continuous and tightly controlled epithelial renewal throughout life (Fig. 1H, I). We detected strong mitotic activity of the *Psc/Su(z)2^DDN^* progenitor cells compared to the control cells as early as three days after induction. Therefore, *Psc/Su(z)2* loss of function generates clusters that are mitotically active and rapidly lose stem cell identity, suggesting that they may have tumor-like characteristics.

As *esg*-positive progenitors include ISCs, EBs and EEps, we asked which cell population was at the origin of the *Psc/Su(z)2* mutant clusters. We used specific drivers allowing the expression of *Psc/Su(z)2* RNAi in each cell type: *Dl-GAL4^ts^* or *esg-GAL4^ts^,Su(H)-GAL80* in ISCs, *Su(H)-GAL4^ts^* in EBs and *Piezo-GAL4^ts^*in EEps. Both ISC drivers showed clusters with small nuclei and Prospero-negative cells, as previously obtained with *esg-GAL4^ts^* (Fig. 1J, K; Fig. S1B, bottom panels). Again, we observed the disappearance of GFP, supporting a loss of driver activity. In contrast, *Psc/Su(z)2* loss of function in EBs does not produce clusters (Fig. 1J, K), whereas in EEps, some guts exhibit the cluster-like phenotype (Fig. 1J, K). This weakly penetrant phenotype observed with the EEp driver is likely due to *Piezo* expression in a subpopulation of Dl-positive cells [37]. Taken together, our results demonstrate that Psc and Su(z)2 act in a cell-autonomous manner in ISCs to prevent the formation of tumor-like clusters.

*Psc* and *Su(z)2* have previously been reported to act redundantly in the *Drosophila* ovary and testis to restrain stem cell proliferation [14, 15]. We used both RNAi and the MARCM system to induce knockdown or knockout of only one of the two genes. Single *Psc* or *Su(z)2* RNAi does not form tumor-like clusters, and the single MARCM clones harbor Dl-positive ISCs and generate differentiated cells (large-nucleated ECs and Pros-expressing EEs) (Fig. S1C-E). These results reveal that *Psc* and *Su(z)2* act redundantly in ISCs, reminiscent of what has been reported in other *Drosophila* organs like the ovary and testis [14, 15].

We then tested by RNAi whether the selective knockdown of the other core PRC1 subunits Sce, Ph, and Pc, as well as of the substoichiometric subunit Scm, could phenocopy the loss of Psc/Su(z)2. We observed that loss of function of the catalytic subunit Sce or the subunit Ph (depletion of both Ph-d and Ph-p) is sufficient to induce the tumor-like phenotype and the disappearance of GFP expression. However, loss of the Pc or substoichiometric Scm subunits does not produce the *Psc/Su(z)2* loss-of-function phenotype (Fig. S2A, B). This is a surprising observation since Pc and Scm, like Psc/Su(z)2, Ph and Sce, are part of the core PRC1. Nevertheless, this result is in accordance with previously published data in imaginal discs showing that the loss of Pc and Scm subunits have no or less severe phenotypes compared to the loss of other core PRC1 subunits [33, 38]. To fully explore the complexity of PRC1 subunit association in this context, we investigated whether a non-canonical PRC1 complex could be involved in ISCs to prevent cluster formation. Therefore, we tested RNAi-induced loss of function of the known non-canonical PRC1 subunits RYBP and Kdm2 and did not observe the formation of cell clusters in any of these conditions (Fig. S2A, B). Overall, our results point to an essential role for the core PRC1 complex, comprising Sce, Ph and Psc/Su(z)2 but independently of Pc, in maintaining ISC identity and ensuring the full differentiation capacity of the stem cell lineage in the adult *Drosophila* intestine. The loss of core PRC1 function in adult ISCs results in the formation of clusters with tumor-like characteristics, which include the loss of cell identity, loss of differentiation and unchecked proliferation.

### *Psc/Su(z)2* loss of function triggers intestinal neoplastic transformation and impairs intestinal response to stress

The tumor-like characteristics of the PRC1 mutant clusters prompted us to investigate whether they were potentially malignant. Neoplastic malignant tumors are expected to be immortal and to acquire metastatic behavior after serial transplantations (allografts) in adult abdomens [39]. We conducted allograft experiments with GFP-positive clusters coming from *esgF/O>Psc/Su(z)2^RNAi^*flies, described above (Fig. S1A), allowing co-expression throughout the process of *Psc/Su(z)2^RNAi^* for cluster formation and of GFP for tracking. We performed two series of allografts, involving 18 transplants from 6 independent *Psc/Su(z)2^RNAi^* adult midguts into the abdomens of healthy adult female hosts. Before killing their hosts, two independent transplants (tumor 1 and tumor 2) were extracted, and portions were re-transplanted into new healthy flies until the fifth generation of transplantation (G1 to G5). The *Psc/Su(z)2^RNAi^*transplants maintained their ability for sustained growth in host flies over the five generations (about 1,5 month) (Fig. 2A). Furthermore, in contrast to control *esgF/O* cells, *esgF/O>Psc/Su(z)2^RNAi^* cells exhibited a propensity to invade the whole host body as early as G1, disseminating from the initial injection site and infiltrating the entire host organism (Fig. 2A). We observed a significant decline in host fly survival that gradually worsened through transplantations, demonstrating that *Psc/Su(z)2*-depleted transplants become more aggressive upon serial transplantation (Fig. 2B-D). These results imply that the PRC1 mutant clusters are neoplastic tumors that exhibit continuous and autonomous growth and are hereafter referred to as tumor-initiating intestinal cells (TIICs).

**Figure 2.**
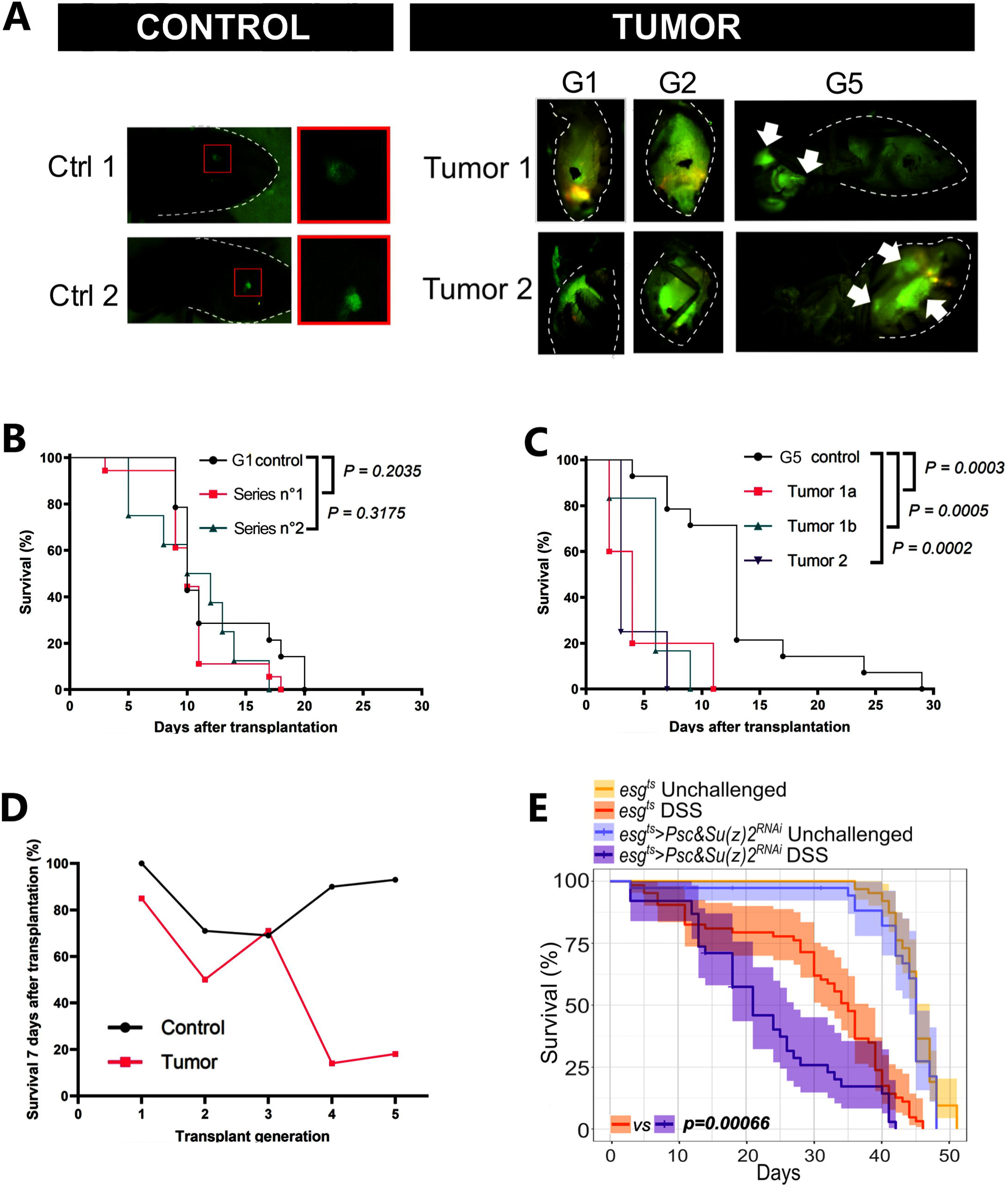
*Psc/Su(z)2* loss of function triggers intestinal neoplastic transformation and impairs intestinal response to stress. **A.** Visualization of the GFP signal (green) after serial transplantation (allograft) of *esgF/O>GFP* intestinal cells into the abdomens of adult female hosts. Left panels: images showing two control flies in which the GFP signal of *esgF/O* midguts (i.e., GFP-positive midguts of siblings coming from the cross allowing the generation of intestinal TIICs) is only visible close to the injection site after the first generation of transplantation. Right panels: Images showing the proliferative and invasive properties of two GFP-labeled *esgF/O>Psc/Su(z)2^RNAi^*intestinal tumors. In the first two generations of transplantation (G1 and G2), the GFP-positive cells already have the ability to disperse in the abdomen. By G5, *Psc/Su(z)2^RNAi^* tumors have acquired the ability to invade different sites, even outside the abdomen region (arrows). The abdomen are limited by white dotted lines. **B**. Survival rate as a percentage of flies after the first generation of transplantation (G1). Two independent transplantation series (series n°1 in pink and series n°2 in green, respectively) were performed and compared to the control flies. **C**. Survival rate (%) after the fifth generation of control flies (G5 control in black; injected with PBS) and *Psc/Su(z)2^RNAi^*tumor-transplanted flies (in pink, green and purple). **D**. Survival (%) of host flies 7 days after control allograft (black) and after allograft of *Psc/Su(z)2^RNAi^* knockdown tumors (red) during 5 rounds of transplantation. In all allograft experiments, significant differences were assessed using the *Log-rank test*. **E**. Survival curves (%) upon DSS challenge of control flies (*esg^ts^*) or flies expressing *Psc/Su(z)2^RNAi^* (*esg^ts^>Psc/Su(z)2^RNAi^*)*. Statistics with Pairwise CoxPH comparisons*.

We next investigated whether TIICs compromise the overall health of the flies in response to extrinsic challenges. To this end, we exposed the flies to dextran sulfate sodium (DSS), which induces gut damages similar to human ulcerative colitis [40]. The survival curves of unchallenged *esg-GFP^ts^*/+ (control) or *esg-GFP^ts^>Psc/Su(z)2^RNAi^* (tumor-bearing) flies did not appear to be significantly different, indicating that TIIC tumors had no direct impact on *Drosophila* lifespan (Fig. 2E). However, *esg-GFP^ts^>Psc/Su(z)2^RNAi^*flies treated with DSS exhibited a significant decrease in survival compared to control flies treated with DSS. These results demonstrate that *Psc/Su(z)2* loss of function impairs intestinal response to colitogenic stress in flies, leading to a shorter lifespan upon challenge. Altogether, these results highlight that core PRC1 is required to maintain ISC identity, thereby ensuring full capacity for gut regeneration. Moreover, preserving normal levels of PRC1 activity is crucial for safeguarding the adult intestine against malignant neoplastic tumors.

Recent studies showed that transient depletion of PRC1 subunits in the larval eye imaginal disc is sufficient to induce a neoplastic transformation, indicating that epigenetic modifications can initiate tumorigenesis independently of genetic mutations [35]. To investigate whether epigenetically induced tumors can also be generated in adult tissues, we used the thermosensitive RNAi system enabling the reversible knockdown of *Psc/Su(z)2* in adult intestinal progenitors, as performed in the eye imaginal disc. We observed that a transient loss of Psc/Su(z)2 for 24 or 48 hours led to the formation of TIIC tumors, with the 48-hour condition achieving an efficacy close to that of continuous 14-day depletion (Fig. S2C, D). Reminiscent of the observations made in the larval eye imaginal disc, these results further demonstrate that a transient loss of the epigenetic modulator PcG genes is also sufficient to induce neoplasia in an adult tissue and that epigenetic alteration can lead to adult cell reprogramming.

### TIIC transcriptomic signatures

To uncover key players leading to the formation of PRC1-dependent intestinal tumors, we compared the transcriptome signatures of control progenitors to the FACS-sorted *Psc/Su(z)2^RNAi^* TIICs. The RNA sequencing data showed that 2488 genes are defined as significantly differentially expressed, with 1268 downregulated genes and 1220 upregulated genes (Fig. 3A; Table S1). The efficacy of the knockdown of *Psc* and *Su(z)2* in TIIC tumors is validated by the reduction of their expression (Fig. 3B). As expected, several canonical Homeotic (*Hox*) PcG targets are derepressed, such as *abd-A*, *Abd-B*, *Antp*, *pb*, *Dfd* and *Scr* (Fig. 3B). Our data confirmed the concomitant loss of *esg* transcription, which is consistent with our previous observation that *esg* expression disappeared in TIICs (Fig. 3B). We also detected decreased expression of several validated ISC markers: *mira*, *insc*, *sox21a* and *zfh2* [41–43]. These data, together with our previous findings (Fig. 1), confirm that TIICs lose stem cell identity.

**Figure 3.**
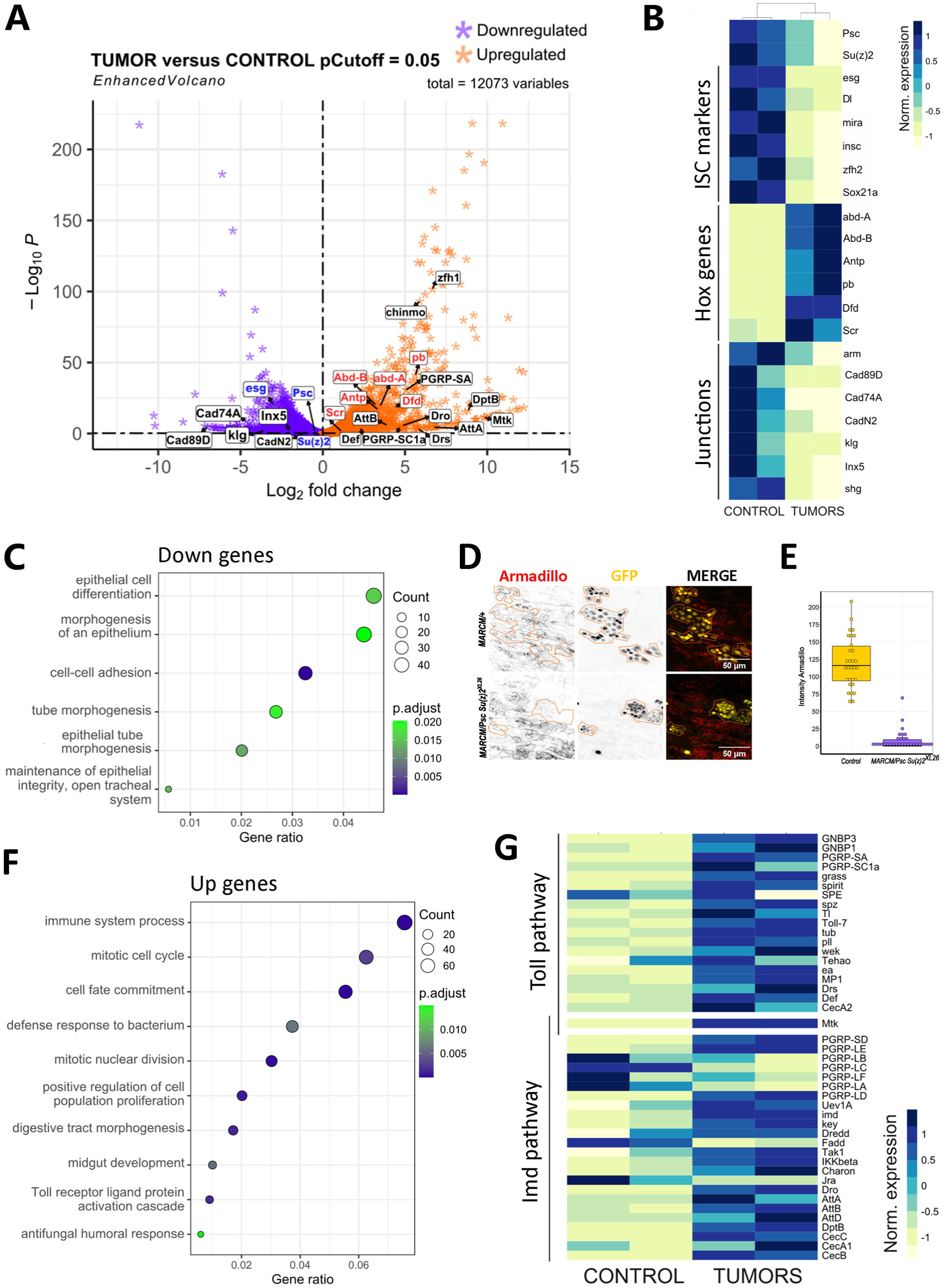
*Psc/Su(z)2* TIIC transcriptomic signature reveals activation of the Toll and Imd pathways. **A**. Volcano plot of differentially expressed genes between *Psc/Su(z)2^RNAi^* tumors and control progenitors. Each star represents a gene whose expression is significantly modified with a log2 fold-change of less than –1 (purple) and more than +1 (orange). A set of candidate genes is indicated (see main text for details). **B**. Heatmap showing normalized expression of key genes in control progenitors and *Psc/Su(z)2^RNAi^*tumors (see main text for details). **C.** Representative Gene Ontology terms enriched in downregulated genes. The full chart is available in Fig. S3. **D.** Control and *Psc/Su(z)2^XL26^* mutant MARCM clones expressing GFP (yellow). The adherent junctions were marked with the anti-Armadillo antibody (red). Clones and clusters are delimited by yellow lines. Scale bar, 50 μm. **E.** Quantification of the intensity of the Armadillo signal in control (orange) and *Psc/Su(z)2^XL26^*mutant (purple) clones. n= 10 guts across 2 independent experiments for a total of 30 measurements for each condition. **F.** Representative Gene Ontology terms enriched in upregulated genes. The full chart is available in Fig. S3. **G.** Heatmap showing normalized expression of genes of the Imd and Toll signaling pathways in control progenitors and *Psc/Su(z)2^RNAi^* tumors.

Gene ontology analysis revealed that downregulated genes are enriched for functions such as “epithelial cell differentiation”, “cell adhesion” and “maintenance of epithelial integrity” (Fig. 3C; Fig. S3; Table S2), including genes encoding for Armadillo/β-Catenin and cadherins (Fig. 3B). Armadillo/β-Catenin, a component of the adherent junctions in the intestinal epithelium that is enriched in ISC [44], shows reduced expression in *Psc/Su(z)2^XL26^*MARCM clones, as evidenced by immunostaining, which validates the loss of epithelial identity in TIIC tumors (Fig. 3D, E). Conversely, the upregulated genes are enriched for ontologies associated with proliferation control like *polo*, *stg* and *Cdk1* (Fig. 3F; Fig. S3; Table S3), which is in accordance with proliferative features and the loss of differentiation that are characteristic of a tumoral phenotype. Data also showed a strong upregulation of genes involved in the immune response against bacteria and fungi (Fig. 3F; Table S3), which was not observed in control midguts dissected under the same conditions. This suggests that the activated immune response is specific to PRC1 downregulation. In particular, several members and target genes of the Toll and Imd signaling pathways showed an increase in transcription in TIICs, including the *PGRP-SA*, *spz*, *Toll*, *Drs* and *Def* for the Toll pathway and *DptB* for the Imd pathway (Fig. 3G; Fig. S3). A recent study used the CUT&Tag methodology to determine the direct targets of the Pc-containing complexes in the whole *Drosophila* gut [45]. Re-analysis of their data indicates that several genes of the Toll/Imd pathways are indeed direct targets of Pc, suggesting that the upregulation of the Toll/Imd pathways in TIIC tumors may be a direct effect of the PRC1 loss (Fig. S3E, F). Interestingly, these pathways were not found to be upregulated upon depletion of PRC1 subunits in the larval eye imaginal disc [35], indicating that this response is specific to the gut.

### Activation of the Toll and Imd pathways supports an antitumoral compensatory mechanism independently of cell death

To assess the potential involvement of immune activation through the Toll and Imd pathways in TIICs, we investigated whether RNAi-induced knockdown of the two downstream NF-κB transcription factors dorsal (Toll pathway) or Relish (Imd pathway) affects the development of TIIC tumors. We observed that the concomitant loss of *Psc/Su(z)2* and *Relish*, or more notably *Psc/Su(z)2* and *dorsal*, induced massive tumors (Fig. 4A, B). In contrast, the loss of *Relish* or *dorsal* alone does not result in tumor formation (Fig. 4A, last two rows). The number of mitotic cells per gut increased significantly in both conditions compared to normal TIIC tumors (Fig. 4C). *Drosophila* Toll and Imd signaling, and their downstream effectors NF-κB dorsal and Relish, have been shown to exert tumor suppressive roles through pro-apoptotic cell death [46–51]. We therefore tested whether a compensatory mechanism takes place in TIICs that reduces tumor expansion by promoting cell death. We expressed p35 or *reaper^RNAi^* in TIICs but observed neither a significant change in the severity of the tumor phenotype compared to *Psc/Su(z)2*-depleted tumors nor in mitotic activity (Fig. 4A-C). In addition, we used the TUNEL assay to detect dying cells, and the results showed no differences in the proportion of cell death between control and TIIC conditions (Fig. 4D, E). Our findings thus indicate that Dorsal of the Toll pathway and Relish of the Imd pathway are both cell-autonomously activated in TIIC tumors to restrict their expansion independently of cell death. However, these compensatory mechanisms are not sufficient to prevent neoplastic tumor formation and progression after transplantation.

**Figure 4.**
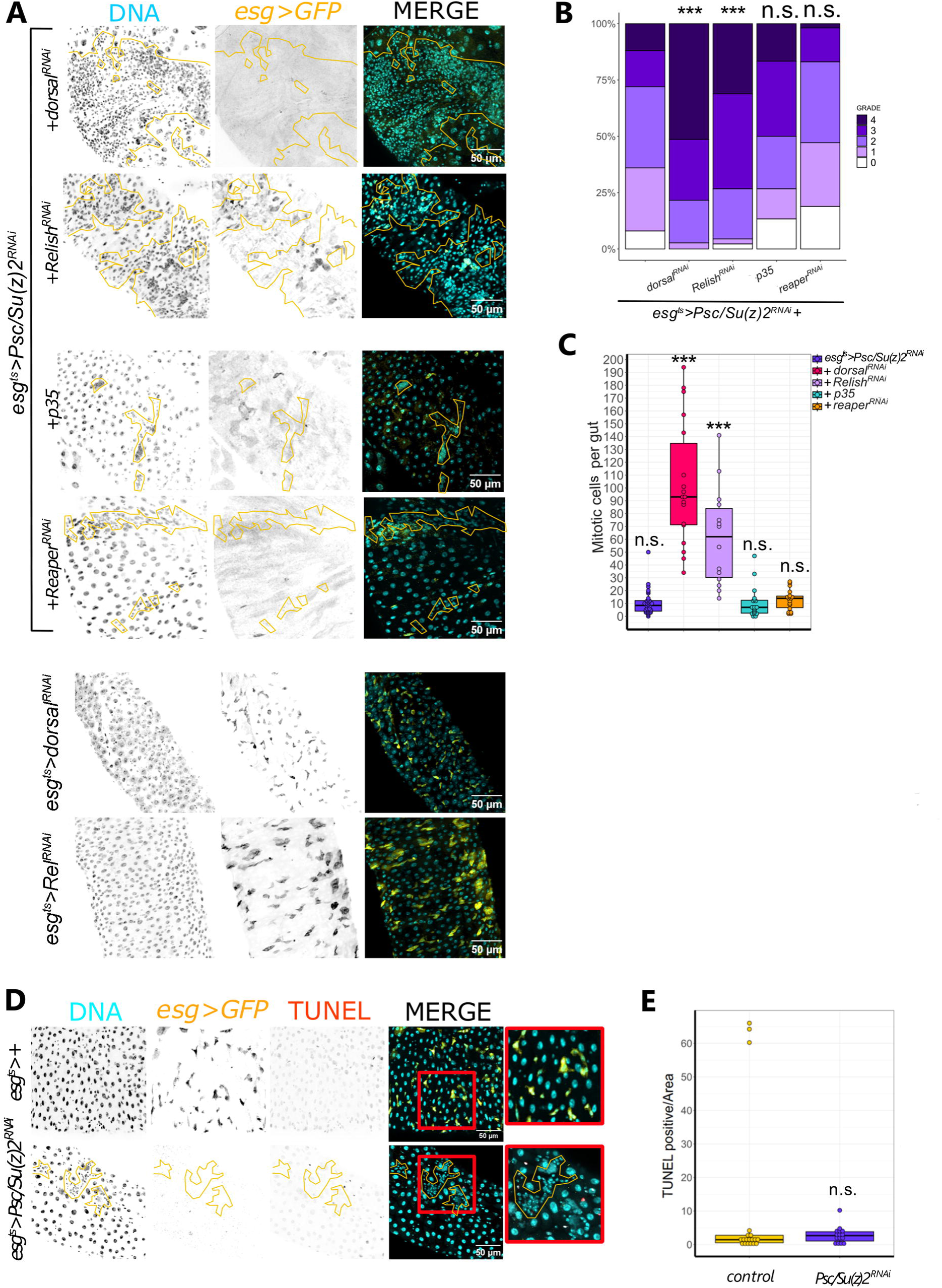
Activation of the Toll and Imd pathways acts as a compensatory mechanism that is independent of cell death. **A**. *esg*-positive cells expressing GFP (yellow) and *Psc/Su(z)2^RNAi^* along with *dorsal^RNAi^, Relish^RNAi^, p35 or reaper^RNAi^*. Tumors are outlined with yellow lines. In contrast, the *esg^ts^>dorsal^RNAi^*and *esg^ts^>Relish^RNAi^* controls showed no tumors (n= 10 for each). **B**. Quantification of the tumor burden following 5 grades of severity (0=no tumor to 4=large tumors) as described in Fig. S4. *P-value 1.6586.10^-5^ and 8.098.10^-5^, respectively, compared to the Psc/Su(z)2^RNAi^ control condition. Statistics were obtained using a contingency table and Chi-sq. tests.* **C**. Quantification of the mitotic cells (PH3-positive) per midgut in the conditions indicated in A. *Anova-Tukey multiple comparisons. P-value <0.0000000 and 0.0000002, respectively, compared to Psc/Su(z)2^RNAi^ control condition*. **D**. *esg*-positive cells expressing GFP only (yellow) or with *Psc/Su(z)2^RNAi^* stained by the TUNEL method (red). Close-up images are also shown (red squares). Tumors are outlined with yellow lines. **E**. Quantification of the TUNEL-positive cells per normalized area in the 2 conditions. *Student T-test shows no significant differences.* DNA (turquoise): stained with DAPI. Scale bar, 50 μm.

### Activation of JAK/STAT signaling sustains TIIC tumor growth

Finally, we went a step further to find the mechanisms that might support tumor formation. Analysis of our RNA sequencing data yielded a marked increase in the expression levels of *chinmo* and *zfh1* (Fig. 5A; Table S1). These two genes have been shown to be direct PcG targets and to promote tumor growth [33, 35, 52–58]. Of particular interest, the mammalian homologue of *zfh1*, *ZEB1*, is an oncogene that has been reported in various human cancers and has the capacity to induce epithelial-to-mesenchymal transition [59]. Interestingly, *chinmo* and *zfh1* have been identified as direct transcriptional targets of the JAK/STAT pathway [55], prompting us to evaluate the role of this pathway in TIIC tumors. Consistent with this, our data showed decreased expression of *ken*, a known negative regulator of the JAK/STAT pathway (Fig. 5A). We first monitored the activity of the pathway using the *10XSTAT92E>GFP* reporter transgene. Tumors obtained either by expression of the DDN forms of Psc/Su(z)2 proteins or by *Psc/Su(z)2* RNAi showed consistent activation of JAK/STAT signaling (Fig. 5B). In the eye imaginal disc, loss of PRC1 drives tumor growth partly in a JAK/STAT-dependent manner through the production of the Upd ligands [60]. We re-analyzed the Pc-profiling CUT&Tag performed in intestinal cells [45], focusing on the JAK/STAT genes found regulated in TIIC tumors. The results indicated that the genes coding for the ligands Upd2 and Upd3 (but not Upd1) can be potentially directly regulated by PRC1 (Fig. S3G). In our RNA sequencing data, however, we did not detect transcriptional upregulation of these ligands, suggesting that JAK/STAT activation in the TIIC tumors does not occur through derepression of the *upd* genes. To directly address whether expression of JAK/STAT pathway ligands is indeed unchanged, we analyzed *upd3-lacZ* expression in guts bearing tumors (*esg^ts^,GFP,upd3-lacZ>Psc-Su(z)2^DDN^*). The results indicate no change in expression of *upd3* in TIICs or adjacent cells compared to control guts, suggesting that JAK/STAT activation is independent of the Upd ligands (Fig. 5C). Finally, an RNAi loss-of-function experiment targeting the JAK/STAT transcription factor Stat92E in *Psc/Su(z)2* TIICs caused a significant decrease in the number and size of the tumors, even though we still observed the loss of the *esg>GFP* marker (Fig. 5D, E). This indicates that the tumor burden of *Psc/Su(z)2* clusters is reduced upon *Stat92E* knockdown. Collectively, these observations indicate that JAK/STAT signaling contributes to the growth of *Psc/Su(z)2*-induced TIIC tumors.

**Figure 5.**
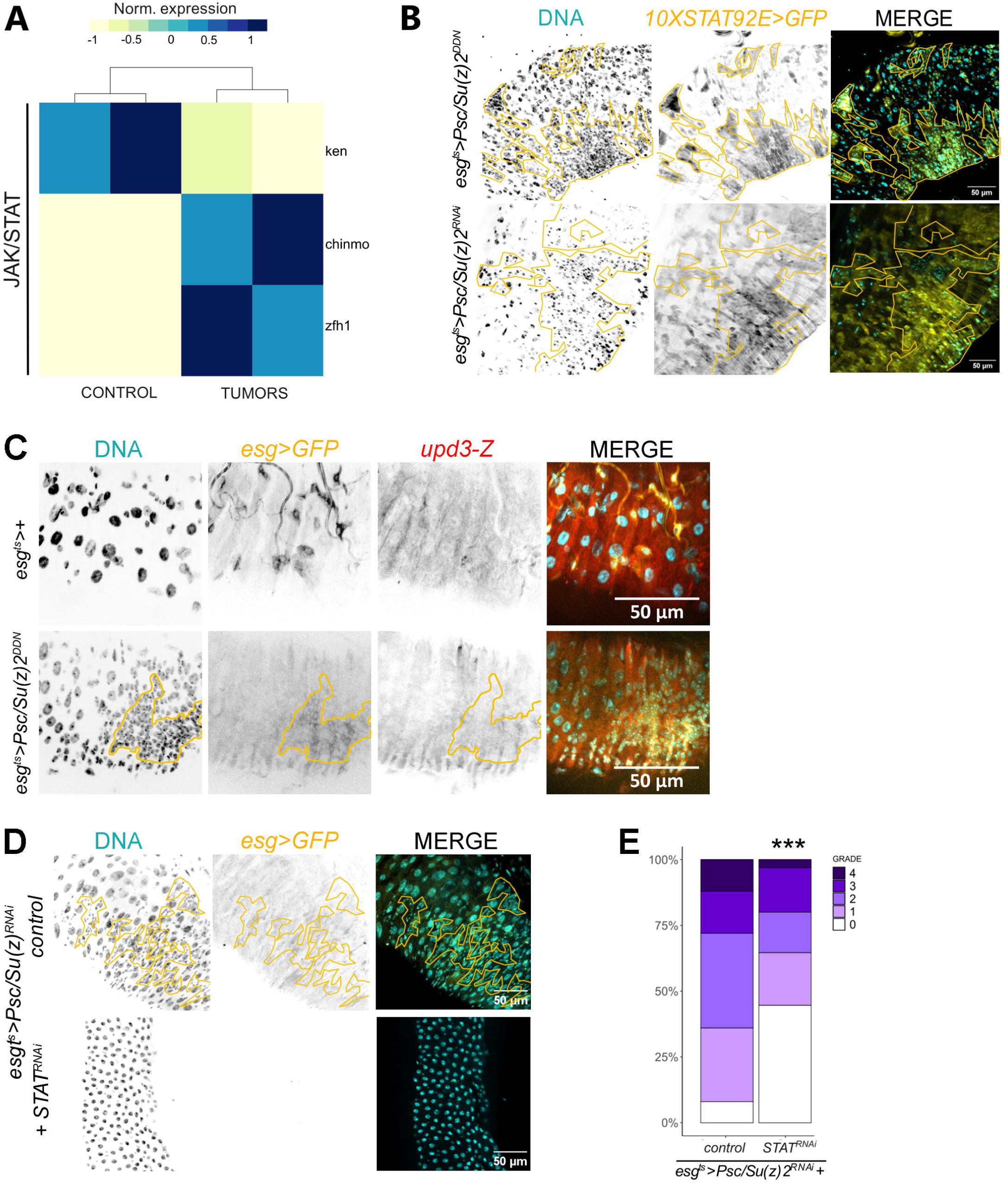
Activation of JAK/STAT signaling supports tumor growth. **A**. Heatmap showing normalized expression of the negative regulator *ken* and the target genes *chinmo* and *zfh1* of the JAK/STAT pathway in control progenitors and *Psc/Su(z)2^RNAi^* tumors. **B**. 10X*STAT92E>GFP* reporter gene expression (yellow) in *esg^tsNP7097^>Psc/Su(z)2^DDN^*(top panels) or *Psc/Su(z)2^RNAi^* tumors (bottom panels). **C**. *upd3-LacZ* reporter gene expression (red) in control guts (*esg^ts^,UAS-GFP>+*; top panels; n= 14) and TIIC-containing guts (*esg^ts^,UAS-GFP>Psc/Su(z)2^DDN^*; bottom panels; n= 23). **D.** *esg*-positive cells expressing GFP (yellow) and *Psc/Su(z)2^RNAi^* only or with *STAT92E^RNAi^*. **E**. Tumor burden in *Psc/Su(z)2^RNAi^* (control) and *Psc/Su(z)2^RNAi^*,*STAT^RNAi^*. *Chi-sq. test p-value= 0.0006242*. Tumor burden was evaluated as described in Fig. 4 and Fig. S4. Tumors are outlined with yellow lines. DNA (turquoise): stained with DAPI. Scale bar, 50 μm.

## DISCUSSION

Our work identified the PRC1 components Psc/Su(z)2, Sce and Ph (members of core PRC1) as crucial tumor suppressors in the adult *Drosophila* intestine. We showed that the loss of function, even transient, of these components in adult ISCs results in the loss of stem cell identity and the formation of JAK/STAT-mediated malignant cell clusters, which we termed tumor-initiating intestinal cells (TIICs) (Fig. 6). Studies in *Drosophila* have provided insight into many mechanisms that lead to tumor initiation and progression, with the intestine being one of the major tissues among adult tumor models [61, 62]. Numerous signaling pathways, including EGFR, Notch, DPP, HIPPO, JNK, Wg/Wnt, JAK/STAT, have been identified as contributing to ISC hyperproliferation and/or tumor formation [63–75]. In these previous studies, tumors manifest hyperplastic characteristics, consistently expressing progenitor, EE and/or EC markers. Importantly, and to the best of our knowledge, the intestinal neoplastic phenotype resulting from the loss of PRC1 function in adult ISCs, characterized by both loss of known intestinal characteristics and continuous growth, has not been previously described in the *Drosophila* midgut and represents a novel tumor model.

**Figure 6.**
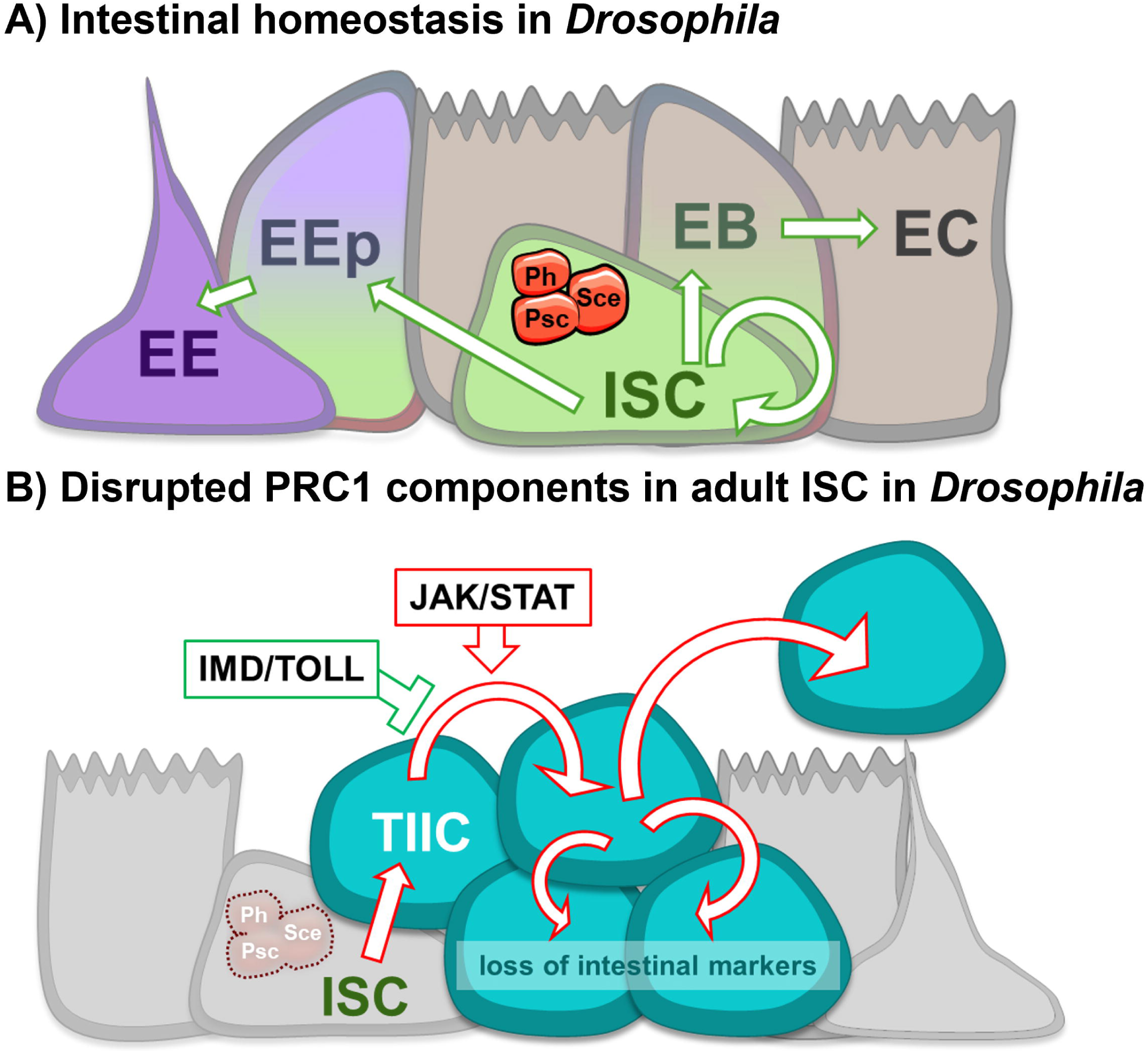
Proposed model of PRC1-mediated maintenance of ISCs in *Drosophila*. **A.** Based on our results, we propose a mechanism whereby the PRC1 components Psc/Su(z)2, Ph and Sce act to control ISC maintenance and proper differentiation mechanisms. **B.** Disruption of Psc/Su(z)2, Ph or Sce in ISCs is sufficient to induce neoplastic cells (TIICs) that lose intestinal identity, thus failing in tissue maintenance and survival under stress. Our results showed that JAK/STAT signaling contributes to TIIC growth, while the Toll/Imd immune pathways limit their expansion independently of cell death.

Epigenetic regulation of adult stem cells is involved in the maintenance of their identity and in their differentiation during homeostasis or aging, but its direct link to tumorigenesis is still little addressed [5, 31, 76]. In the mouse intestine, PRC1 is essential for maintaining the pool of LGR5+ ISCs by supporting Wnt/β-Catenin signaling [77]. PRC1 inactivation leads to a general loss of intestinal lineage identity without the acquisition of any defined differentiation program, reminiscent of what we have observed in the *Drosophila* gut. However, unlike our findings, the loss of PRC1 activity did not result in tumor formation but rather in a progressive reduction of the ISC pool, indicating different outcomes in the mouse and *Drosophila* guts. In *Drosophila*, the Pc subunit is responsible for the age-related deregulation of the ISC lineage, promoting the EE cell fate during aging [32]. Consistent with this specific role, we did not observe intestinal tumor formation upon Pc depletion. This suggests that the Pc subunit is not essential for the silencing function of PRC1, although it potentiates silencing via binding to the H3K27me3 mark. Previous studies have shown that the *Drosophila Psc* and *Su(z)2* genes play redundant roles, for instance in maintaining the somatic stem cell fate of ovarian follicular stem cells in females and of cyst stem cells in males [14, 15]. Like in the intestine, loss of *Psc*/*Su(z)2* function induced the formation of tumorigenic cells. However, in the ovary, this process occurred independently of the other PRC1 subunits and involved sustained activation of the Wg/Wnt signaling pathway [14]. In the testis, only Pc among the PRC1 subunits has been investigated, so there is no evidence for the contribution of the other subunits as a complex [15]. In addition, the tumor clusters only partially lose the identity of the cyst somatic stem cells. Therefore, the mechanisms of action and the sub-complexes of PRC1 involved in stem cell maintenance may vary depending on the adult tissue. Interestingly, a recent study demonstrated that the loss of PRC2 subunits in ISCs results in progenitor depletion due to precocious differentiation of enteroblasts into enterocytes [78]. This phenotype, which is completely different from that observed upon loss of PRC1, indicates that PRC1 and PRC2 complexes act independently and possess distinct functions in ISCs, similar to other *Drosophila* epithelial tissues where a functional uncoupling between PRC1 and PRC2 has been described [79, 80]. Several vertebrate orthologues (PCGF1 to PCGF6) of the *Drosophila Psc* and *Su(z)2* genes have been identified [6]. Some of these orthologues have been directly implicated in cancer [9]. For instance, the *Bmi-1/PCGF4* gene has been the subject of numerous studies for its proto-oncogenic properties, in addition to its role in regulating the self-renewal and differentiation of normal adult stem cells in various tissues [81]. This gene is upregulated in many cancers, including colon cancer, and has emerged as a promising therapeutic target. In particular, *Bmi-1/PCGF4* is important for self-renewal of cancer stem cells and may be involved in tumor initiation [82]. In contrast, *Mel-18/PCGF2* has an antagonistic function towards *Bmi-1* and acts as a tumor suppressor [12, 13]. These data suggest that *Psc* and *Su(z)2* are the functional orthologues of *Mel-18*.

NF-κB signaling is known to exert either tumor-promoting or tumor-suppressing activities depending on the cellular context [83, 84]. In *Drosophila* larvae, two recent studies using oncogenic Ras^v12^-derived tumor models of the eye-antennal disc showed that Toll-NF-κB signaling promotes tumor growth by blocking cell death and differentiation [85, 86]. In contrast, we showed that the Toll pathway and the second Imd immune pathway restrict tumor growth independently of cell death, revealing opposite roles for NF-κB signaling in larval disc tumors and adult TIICs. In vertebrates, a study reported that an increased expression of PcG proteins in cutaneous squamous cell carcinoma represses the NF-κB signaling pathway to facilitate tumor immune escape during metastasis [87]. In *Drosophila*, the Toll and Imd pathways have also been involved in tumor suppression, but no link with PcG proteins has yet been found [46–51]. Toll signaling is activated in larval wing imaginal discs by the ligand Spätzle, which is locally secreted by Myc-overexpressing cells to induce apoptosis in flanking cells [46]. These pathways can also exert their non-cell autonomous tumor suppressor functions through the activation of circulating hemocytes or the systemic production of antimicrobial peptides (AMPs) [47, 50, 88, 89]. In contrast, we have shown that these pathways act cell autonomously in the gut, as their inactivation within tumor cells is sufficient to increase neoplasia. The question then arises as to how Toll and Imd signaling restrict intestinal tumor growth in adult tissues like the intestine. One possibility involves the production of AMPs, as seen in larval imaginal discs and lymph glands [47, 50]. However, these studies have shown that AMPs control tumorigenesis by killing tumor cells in an apoptosis-dependent manner. Since we did not detect cell death in intestinal tumors, it is likely that a different mechanism of tumor suppression occurs in the adult gut.

JAK/STAT signaling has been involved in various types of cancer in both *Drosophila* and humans [90, 91]. In *Drosophila*, sustained JAK/STAT activation during development promotes epithelial tumors in imaginal discs, as well as hematopoietic and melanotic tumors, whereas little is known about its role in adult tissues. While our findings strongly link PRC1 loss to the emergence of malignant JAK/STAT-driven TIICs, technical limitations hampered our ability to determine which genes are directly repressed by PRC1 to maintain stem cell identity and prevent neoplasia. Prior ATAC-seq analysis in a related context, i.e. sorted ISCs with *Pc* knockdown during aging, revealed an unexpected very modest decrease in global chromatin accessibility, suggesting that PRC1 may also repress transcription through mechanisms not strictly dependent on promoter accessibility [32]. This complicates the interpretation of chromatin state as a readout for PRC1 activity and suggests that epigenetic repression may involve alternative or indirect pathways. Our efforts to perform CUT&RUN experiments for PRC1-associated histone marks (H3K27me3 and H2AK118Ub) on FACS-sorted ISCs were unsuccessful, largely due to technical constraints including limited starting material and poor signal recovery from rare cell populations. These challenges underscore a broader methodological gap in epigenomic profiling of *Drosophila* ISCs and highlight the need for improved techniques designed for low-input samples. Future technical innovations will be required to fully elucidate the role of PRC1 in stem cell regulation and cancer suppression in the *Drosophila* gut.

Our work shows that, in adult tissue, modification in the epigenetic regulation of gene transcription by PRC1 is sufficient to promote tumorigenesis. This finding aligns with previous research in the developing eye imaginal disc, where PRC1-dependent tumor growth also involves activation of the JAK/STAT pathway [35]. However, our study also reveals tissue-specific mechanisms, since the Toll and Imd pathways are uniquely activated in TIIC tumors to limit their progression. Interestingly, the gut constitutes an important interface with microbes, where the immune system is continuously solicited through interactions with the microbiota and encounters with pathogens. Our findings underscore how the gut’s primary role in protecting the organism from external aggressions extends beyond pathogen defense to also mitigate tumorigenesis. This highlights a fascinating interplay between tissue function and disease modulation. Our study emphasizes the need to further explore epigenetic regulatory mechanisms in cancer research, and understanding these processes could open new avenues for therapeutic strategies targeting epigenetic modifications.

## MATERIALS & METHODS

### Reagents and biological resources

#### Antibodies

Anti-Armadillo antibody was deposited to the DSHB by Wieschaus, E. (DSHB #N2 7A1 ARMADILLO-s)

Anti-Prospero was deposited to the DSHB by Doe, C.Q. (DSHB #MR1A)

Anti-Delta was deposited to the DSHB by Artavanis-Tsakonas, S. (DSHB #C594.9B)

Anti-Phospho-Histone 3 (Cell signaling #9701S)

Anti-β-Galactosidase (Gentex #GTX77365)

#### Chemicals and consumables

PBS-10X (Euromedex #ET330)

Triton– X-100 (Sigma #T9284)

16% Formaldehyde solution Methanol-free (ThermoScientific #28900)

Hoechst (Molecular Probes H-3569; 1/1000)

Fluoroshield DAPI medium (SIGMA #F6057-20mL)

Fluoromount Aqueous mounting medium (SIGMA #F4680-25mL)

Elastase from porcine pancreas (SIGMA #E7885-5MG)

PluriStrainer Mini 40 µm (PluriSelect #43-100040-40)

Glycerol (SIGMA #G5516)

In situ Cell Death Detection Kit (ROCHE #12156792910)

Dextran sulfate sodium salt (SIGMA #42867-25G)

#### *Drosophila* strains

Wild-type *Canton S* (RRID:BDSC_64349)

*w; esg-GAL4^NP5130^ UAS-GFP; tubGAL80ts* (gift from Yiorgos Apidianakis)

*w; esg-GAL4^NP7097^; tubGAL80ts* (gift from Nic Tapon)

*esg-GAL4,UAS-2xEYFP/CyO; Su(H)GBE-GAL80,tub-GAL80ts* (gift from Bruce Edgar)

*esg-GAL4, UAS-2EYFP, Rel^RNAi^/Cyo; Su(H)-GAL80, tubGAL80ts/MKRS* [92]

*esg-GAL4, tub-GAL80ts, UAS-GFP; UAS-Flp, Act > CD2 > GAL4* (*esgF/O*) (gift from Bruno Lemaitre)

*esg-Gal4, tub-Gal80ts, UAS-GFP/CyO; upd3-lacZ/TM6* (gift from Michael Boutros)

*w; tub-GAL80ts; Dl-GAL4 UAS-GFP/TM6b* [93]

*w; Su(H)GBE-GAL4/SM6β; tub-GAL80ts UAS-GFP/TM6b* [93]

*w; Su(H)GBE-LacZ* (gift from Julien Colombani)

*w; UAS-GFP, Piezo-GAL4[KI]; tubP-GAL80ts* (RRID:BDSC_78337)

*w; UAS-Psc.N1.Myc,UAS-Su(z)2.N1.Myc/CyO* (*Psc/Su(z)2^DDN^*) (RRID:BDSC_68225)

*UAS-RNAi Psc* (RRID:BDSC_38261)

*UAS-RNAi Su(z)2* (RRID:BDSC_33403)

*UAS-RNAi Psc/Su(z)2 #44* & #78 (*Psc/Su(z)2^RNAi^*; recombinant *Psc^RNAi^* from RRID:BDSC_38261 and *Su(z)2^RNAi^* from VDRC #100096) [35]

*UAS-RNAi Ph* (VDRC #50028 & #10679)

*UAS-RNAi Pc* (RRID:BDSC_33622)

*UAS-RNAi Sce* (RRID:BDSC_31612 & RRID:BDSC_35446)

*UAS-RNAi Scm* (RRID:BDSC_55278) *UAS-RNAi Kdm2* (RRID:BDSC_33699)

*UAS-RNAi RYBP* (RRID:BDSC_33974) *UAS-RNAi STAT92E* (VDRC #106980)

*UAS-RNAi reaper* (VDRC #12045)

*UAS-RNAi dorsal* (RRID:BDSC_27650)

*UAS-RNAi Relish* (VDRC #49413)

*UAS-p35* (gift from Tony Ip)

*hs-FLP, tub-GAL4, UAS-GFP/FM; FRT42D, tub-GAL80/CyO* (*MARCM FRT42D*) [14]

*w; FRT42D, Psc-Su(z)2 XL26 / Cyo* [14]

*w; FRT42D Psc e24/CyO* [14]

*y w FRT42D Su(z)2[1.b7]/T(23)TSTL14, SM5: TM6B, Tb[1]* [14]

*10Xstat92E-GFP* (RRID:BDSC_26198)

*Delta.LacZ* (RRID:BDSC_11651)

*w; His2Av-mRFP1* (RRID:BDSC_23650)

#### Softwares and algorithms

FIJI/ImageJ (https://imagej.net/software/fiji/); R (The R project; https://www.r-project.org/); Krita (https://krita.org/fr).

### Experimental methods

#### Drosophila genetics

All *Drosophila* stocks were reared at 25°C on standard medium (0.8% agar, 2.5% sugar, 8% corn flour, 2% yeast) with a 12h light/12h dark cycle. For transgene expression using the GAL4/GAL80^ts^ system, mating experiments were conducted at the permissive temperature for GAL80 protein (18°C). Female flies with the desired genotype were collected within 4 days after hatching and allowed to age for 3-7 days before temperature shift to 29°C to induce transgene expression. Adult midguts were dissected after 14 days of induction or otherwise as indicated. For the MARCM system, 3-7 day old female flies were heat shocked three times at 37°C for 45 min within a period of 1.5 days and maintained at 25°C for 14 days prior to midgut dissection. The transient expression experiment was carried out using the thermosensitive system described previously [35] with the *esg-GAL4^ts^,UAS-GFP* and *Psc/Su(z)2^RNAi^* fly strains. Briefly, transgene expression was induced at 29°C for 24h or 48h before the system was switched off at 18°C for 13 or 12 days, respectively, allowing potential tumors to develop and be compared to a 14-day induction control.

#### Survival experiment

DSS was prepared by dissolving DSS in 1X-PBS/5% sucrose solution, filter sterilized, and stored at 4°C for up to two weeks. For DSS survival, 20 virgin females per vial were reared at 29°C and transferred three times per week to fresh vials containing 5% DSS/5% sucrose or 5% sucrose alone. Dead flies were counted every 1 to 3 days.

#### Gut dissection and staining

Dissections and immunostainings were performed as described [93] using mouse anti-Prospero at 1:200, rabbit anti-PH3 at 1:500, mouse anti-Armadillo at 1:25 and chicken Anti-β-Galactosidase at 1:1000 for the primary antibodies. For Delta staining, we used the previously described protocol [41] using mouse anti-Delta at 1:2500. Samples were mounted with DAPI-containing Fluoroshield medium. Cell death was detected using the TMR red In Situ Cell Death Detection Kit (Roche) following the kit’s standard staining protocol. Briefly, guts were washed in PBS after secondary antibody, then stained with 100uL of TUNEL solution for 1hr at 37°C and washed twice with PBS before mounting.

#### Allograft assay

Allografts were performed as previously described [39]. Briefly, *UAS-RNAi Psc/Su(z)2* #44 female flies were crossed with *esgF/O* males at 18°C. After 3 weeks, 0-3 day old female flies were collected along with a few males, and the intestines were allowed to mature for 3 or 4 days at 18°C. The flies were then transferred to 29°C for 14 days for tumor formation. Tumors were positively labeled with GFP. Only female midguts were dissected, cut into pieces and injected into the abdomens of RFP+ adult female hosts (genotype RRID:BDSC_23650). Flies were monitored every two days, and tumors were dissected and re-injected when the host abdomen was fully GFP. For the first generation of transplantation (G1), the midguts of esgF/O>GFP-positive *CyO* siblings were injected as controls. For subsequent transplantations (G2 to G5), controls were injected with PBS.

#### Cell sorting and RNA sequencing

Female flies with the desired genotype (*esg-GAL4^ts^,UAS-GFP/+* to express GFP in control progenitors; *esg^F/O^/Psc-Su(z)2^RNAi#44^*to induce constitutive GFP expression in TIICs) were collected within 4 days after hatching and allowed to age for 3-7 days at 18°C before temperature shift to 29°C to induce transgene expression for 14 days. 100-120 guts per condition were dissected and placed in a cold vial containing 0.4mL of 1X-PBS. The PBS was discarded and the guts were placed in 10µL of 10mg/mL elastase and 90µL of Tris-HCl pH=8.8. Vials were incubated at 27°C, 60 rpm for 30 minutes with mixing every 10 minutes. Vials were centrifuged at 300g for 20 minutes at 4°C and the pellet was washed in 1X-PBS before reiterating the centrifugation step. Excess PBS was discarded and the pellet was resuspended in 500µL PBS and filtered with PluriStrainer Mini 40 µm filters before GFP-positive cells were sorted by cytometry. Cells were then snap frozen at –80°C and sent to Azenta Life Sciences for mRNA extraction, library preparation (ultra-low input) and sequencing (Illumina HiSeq 2×150bp).

#### RNA-sequencing analysis

Raw data were aligned to the *D. melanogaster* Dm6 (dmel_r6.32) genome assembly. Differential expression analysis of the data was performed using the DESeq2 R package v.1.40.2 (design= ∼ condition). Genes were considered differentially expressed if Padj < 0.05 and |log2fold2 fold change | >1. Principal component analysis (PCA) was performed for quality assessment (Fig. S3A). The Volcano plot was generated using the EnhancedVolcano R library v.1.18.0. Gene Ontology analysis was performed on differentially expressed genes using the *enrichGO* function from the clusterProfiler R package v. 4.8.3, and plots were produced using the ggplot2 R library v. 3.5.1.

#### Pc-profiling CUT&Tag

We re-analyzed the Pc-profiling CUT&Tag dataset recently published [45] as follows. FASTQ data were downloaded from SRA run selector (GSE291173). Reads were aligned to the *D. melanogaster* reference genome dm6 using Bowtie 2 (v 2.4.2). Then, samtools (v1.9) was used to filter out low-quality reads (command ‘samtools view –b –q 30’) and sambamba (v 1.0.0) was used to sort (command ‘sambamba sort’), deduplicate and index BAM files (‘sambamba markdup –remove-duplicates’) with default parameters. For visualization, reads per kilobase per million mapped reads (RPKM)-normalized 27igwig binary files were generated using the bamCoverage function from deepTools2 (version 3.5.5). Peak calling was performed with each replicate as a separate input file using MACS2 with the following parameters: –g dm –f BAMPE –q 0.005. Only peaks from the merged replicates, detected in the four replicates and using merged replicates, were retained for further analysis. Gene Ontology analysis was performed on differentially expressed genes using the enrichGO function from the clusterProfiler R package v. 4.8.3, and plots were produced using the ggplot2 R library v. 3.5.1.

### Imaging, quantification and statistics

#### Image Capture and Processing

All images and data presented were acquired from the R4-R5 posterior midgut using a Zeiss AxioImager Z1 microscope (with Apotome 2 module) at 20x objective, or a spinning disk confocal microscope (WaveFX) at 20x objective. FIJI/ImageJ and Krita software were used for image analysis.

#### Measurements

Fluorescence quantification of Armadillo staining was performed using FIJI/ImageJ. A total of 30 regions of interest (ROIs) per condition were analyzed, derived from n= 10 guts across 2 independent experiments (1–9 ROIs per gut). Two conditions were compared: (i) MARCM mutant clones and (ii) MARCM wild-type clones, which were selected manually based on clone morphology and GFP expression. For each ROI, mean fluorescence intensity was measured using FIJI/ImageJ. Mitotic indexes were determined by manually counting PH3-positive cells in the whole gut. The presence of differentiated cells in clones was evaluated by counting the number of clones containing at least one Prospero-positive cell (from the enteroendocrine lineage) and/or one large nucleus specific for polyploid enterocytes (from the absorptive lineage). In the same way, the presence of stem cells in clones was evaluated by counting the number of clones containing at least one Delta-positive cell. TUNEL-positive cells were counted in FIJI and normalized to area. Tumor burden was determined in a blinded analysis on FIJI following the chart in Figure S4.

#### Statistics

For all experiments, flies were first selected by genotype and then randomly chosen for experimental analysis. All experiments were independently repeated at least 2-7 times. All statistical tests were performed with R software. Box-plots are defined as follows: center line, median; box limits, upper and lower quartiles; whiskers, 1.5x interquartile range; points, outliers. The statistical tests (all two-sided) used to determine the significance, the total number of guts analyzed (“*n*”), and the exact p-value are indicated in the legend of each figure.

#### Ethics statement

We have received authorization from the French ministry of higher education, research and innovation “for the contained use of genetically modified organisms for research, development or educational purposes”, which allows us to handle genetically modified *Drosophila* (DUO number 12508).

## DATA AVAILABILITY

All data supporting the findings of this study are available in the main text or the supplementary materials. The RNA sequencing datasets generated and analyzed during the current study are available in the European Nucleotide Archive under the accession number PRJEB87242 (https://www.ebi.ac.uk/ena/browser/view/PRJEB87242).

## ACKNOWLEDGEMENTS

We wish to thank Yiorgos Apidianakis, Frédéric Bantignies, Michael Boutros, Julien Colombani, Bruce Edgar, Christian Ghiglione, Tony Ip, Bruno Lemaitre, Nic Tapon, Rongwen Xi, the Bloomington Drosophila Stock Center (BDSC; NIH P40OD018537), the Vienna Drosophila Resource Center (VDRC, www.vdrc.at) and the Developmental Studies Hybridoma Bank (DSHB, created by the NICHD of the NIH and maintained at the University of Iowa, Department of Biology, Iowa City, IA 52242) for fly stocks and reagents. We thank Giacomo Cavalli for very insightful comments on the manuscript. We acknowledge Olivier Pierre from the imagery platform of our institute Sophia Agrobiotech for his assistance, and the Cell Imaging Center of the Faculty of Medicine & Dentistry of the University of Alberta. We also acknowledge Julie Cazareth from the flow cytometry and cell sorting platform of the Institute of Molecular and Cellular Pharmacology at Sophia Antipolis for her help with FACS sorting. We are grateful to the bioinformatics and genomics platform, BIG Sophia Antipolis (ISC plantBIOs, https://doi.org/10.15454/qyey-ar89), for computing and storage resources. We thank all the members of the BES (“*Bacillus*, Environnement et Santé”) lab for fruitful discussions.

## AUTHOR CONTRIBUTIONS

Conceptualization: A.J., R.L., A.G., A-M.M., R.R.

Methodology: A.J., A-M.P., A-M.M., R.R.

Validation: A.J., A-M.P., C.R., A-M.M., R.R.

Formal Analysis: A.J., A-M.P., C.R., A-M.M., R.R.

Investigation: A.J., J.S., A-M.M., R.R.

Resources: A.J., A-M.P., C.R., A.G., E.F., A-M.M., R.R.

Data Curation: A.J., A-M.P., C.R., R.R.

Writing – Original Draft: A.J., R.R.

Writing – Review & Editing: A-M.P., C.R., J.S., A.G., E.F., A-M.M.

Visualization: A.J., A-M.P., A-M.M.

Supervision: R.R.

Project Administration: R.R.

Funding Acquisition: A.G., E.F., A-M.M., R.R.

## SUPPORTING INFORMATION CAPTIONS

**Figure S1.** **A**. *esg*-positive cells expressing GFP (yellow) alone (control; upper panels) or together with *Psc/Su(z)2^RNAi^*^#78^ (second row panels) or double dominant negative forms of Psc and Su(z)2 (third row panels). *esg*-positive daughter cells (*esg^ts^ F/O*) expressing GFP together with *Psc/Su(z)2^RNAi^*^#44^ (used in main figures, fourth row panels) or double dominant negative forms of Psc and Su(z)2 (fifth row panels). The differentiated EE were labeled with the anti-Prospero antibody (red). **B**. Same three top panels as in D with the ISCs labeled with anti-Delta (Magenta). *Psc/Su(z)2^XL26^* mutant MARCM clones expressing GFP (yellow) together with the ISC reporter gene *Delta.LacZ* (Magenta, fourth row panels). *esg-Gal4^ts^,Su(H)-*Gal80 (ISC^ts^) expressing GFP (yellow) together with *Psc/Su(z)2^RNAi^*^#44^ (bottom panels). **C**. *Psc^e24^* (single *Psc* mutant; top panels) or *Su(z)2^1b7^* (single *Su(z)2* mutant; second row panels) MARCM clones expressing GFP (yellow). *esg*-positive cells expressing GFP (yellow) together with *Psc^RNAi^* (alone; third row panels) or *Su(z)2^RNAi^* (alone; bottom panels). The differentiated EE were marked with the anti-Prospero antibody (red). **D**. *Psc^e24^* (single *Psc* mutant; top panels) or *Su(z)2^1b7^* (single *Su(z)2* mutant; second row panels) MARCM clones expressing GFP (yellow) with ISCs labeled with the anti-Delta antibody (red). **E**. Quantification of guts with tumor phenotype in %. *Anova-Tukey multiple comparisons. One dot represents one replicate of n= 8-15 guts each*. Tumors are outlined with yellow lines. DNA (turquoise): stained with DAPI. Scale bar, 50 μm.

**Figure S2.** **A**. RNAi loss of function targeting the PRC1 subunits Sce, Ph, Pc, Scm, KDM2 and RYBP in *esg*-positive cells expressing GFP (yellow). **B**. Quantification of guts with tumor phenotype in %. **C.** *esg-*driven expression of GFP and *Psc/Su(z)2^RNAi^* for 14d continuously or transiently for 24 or 48h followed by a period of 13 days or 12 days, respectively, before dissection (to reach a total of 14d). **D.** Quantification of the number of cells per tumor in each condition. *Anova-Tukey multiple comparisons. One dot represents one replicate of n= 8-12 guts each*. Tumors are outlined with red lines. DNA: stained with DAPI. Scale bar, 50 μm or 100 µm as indicated.

**Figure S3.** **A**. Principal component analysis (PCA) of normalized RNA-seq read counts for tumor and control conditions. Each dot corresponds to one biological replicate. Similarity between samples from one condition is reflected in their close distance. **B.C**. Gene Ontology terms enriched in downregulated (B) and upregulated (C) genes of the RNA sequencing data. **D**. Toll (top) and Imd (bottom) pathways in *Drosophila*, from flybase data. **E.** Selected Gene Ontology terms enriched in the Pc-profiling CUT & Tag data recently published [45], after our own re-analysis. **F.** Genes of the Toll/Imd pathways that are direct targets of Pc (CUT & Tag; in orange) in association with the heatmap of their transcriptional regulation in TIIC tumors (RNA sequencing). **G.** Genes of the JAK/STAT pathway that are direct targets of Pc (CUT & Tag; in orange) in association with the heatmap of their transcriptional regulation in TIIC tumors (RNA sequencing). 1D = 1 day-old flies (young); 15D = 15 day-old flies (middle-aged).

**Figure S4.** Tumor burden reference table. Following the table below, tumor burden was determined by analyzing images in FIJI, with image names concealed to ensure a blind assessment. Tumor grades were classified as follows: *Grade I: rare clusters of few cells; Grade II: several clusters of few cells; Grade III: medium size clusters; Grade IV: massive clusters*.

**Table S1.** TIIC RNA sequencing data. Complete list of *Drosophila* genes with their Log2 fold change (Log2FC_TUM) and adjusted p-value (padj_TUM) derived from the RNA sequencing data

**Table S2.** Gene ontology of the downregulated TIIC genes. List of the GO terms (with a p-value < 0.05) corresponding to the downregulated genes identified from the RNA sequencing data.

**Table S3.** Gene ontology of the upregulated TIIC genes. List of the GO terms (with a p-value < 0.05) corresponding to the upregulated genes identified from the RNA sequencing data.

