## Supplementary figures and images for "A tumor-suppressive role of the PRC1 Polycomb epigenetic complex in the maintenance of adult *Drosophila* intestinal stem cell identity"

### Figure S1

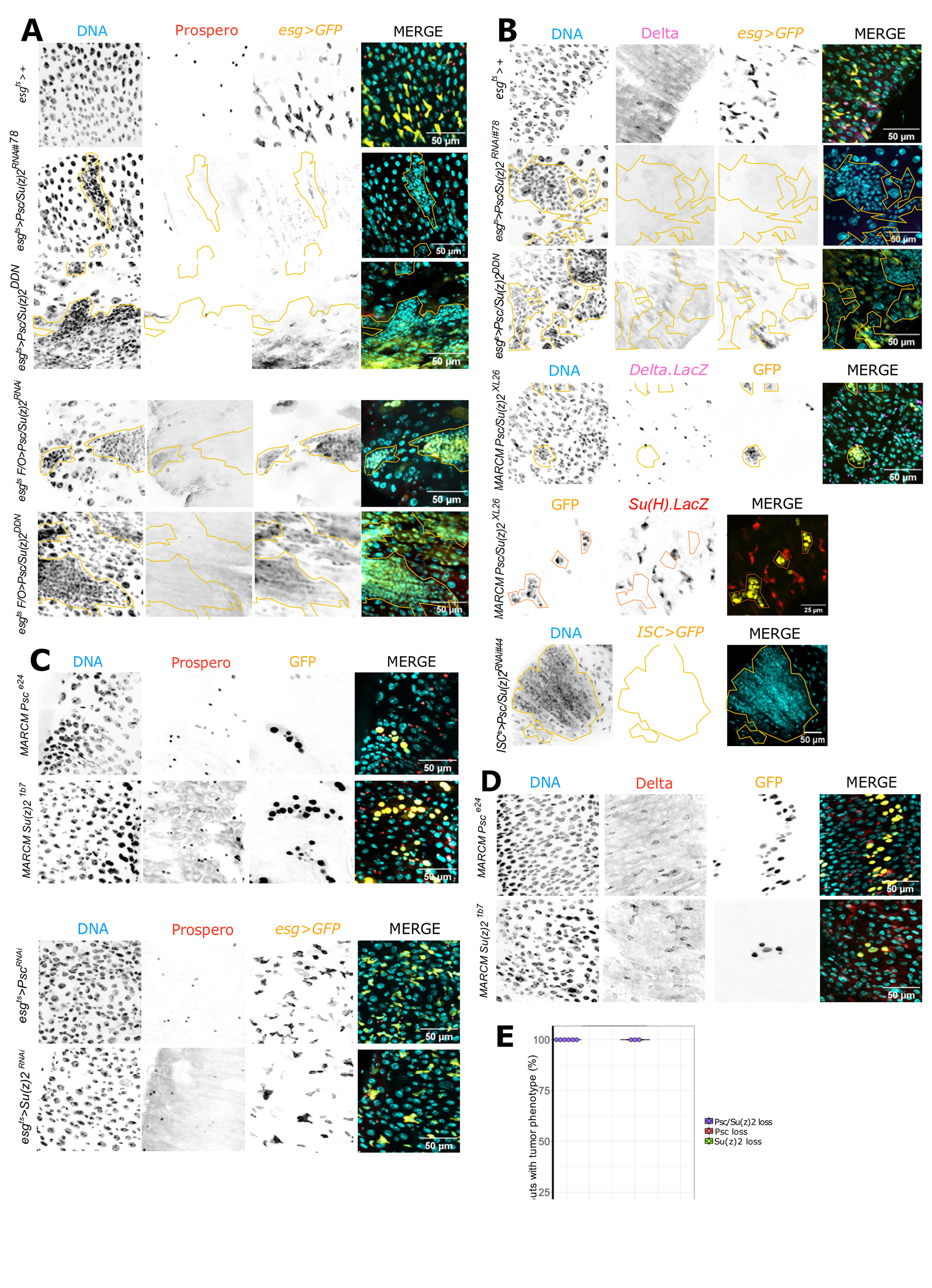

### Figure S2

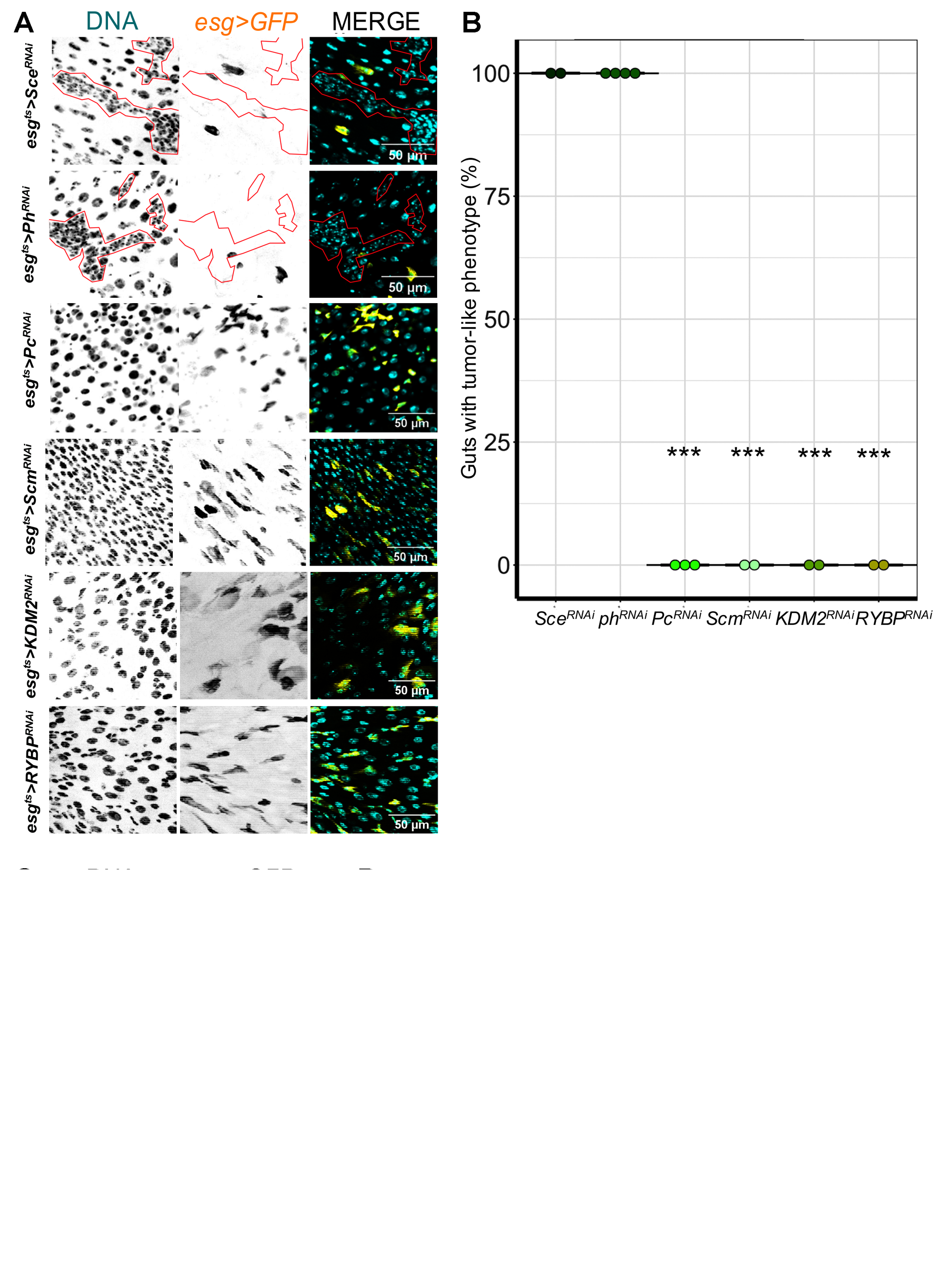

### Figure S3

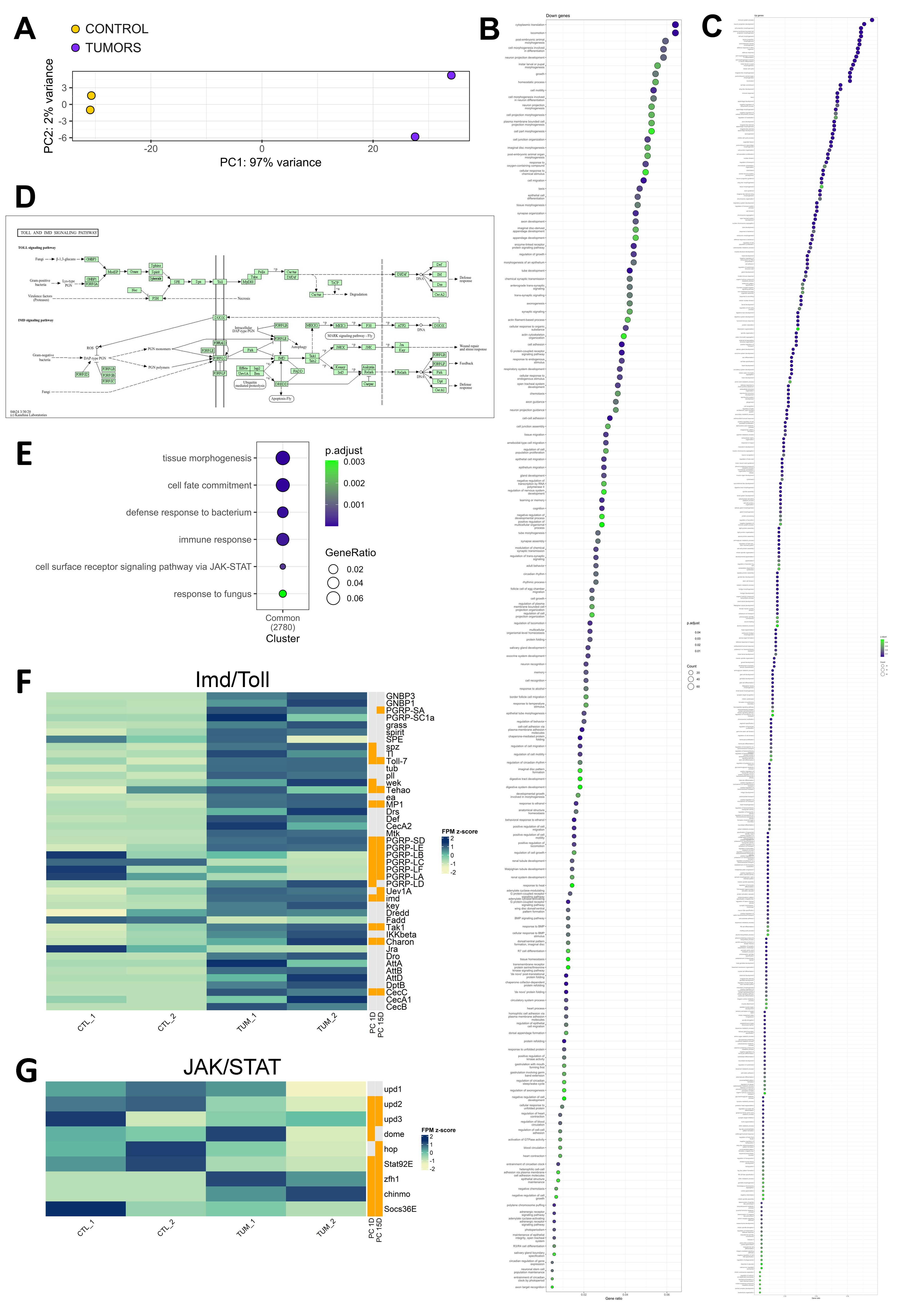

### Figure S4

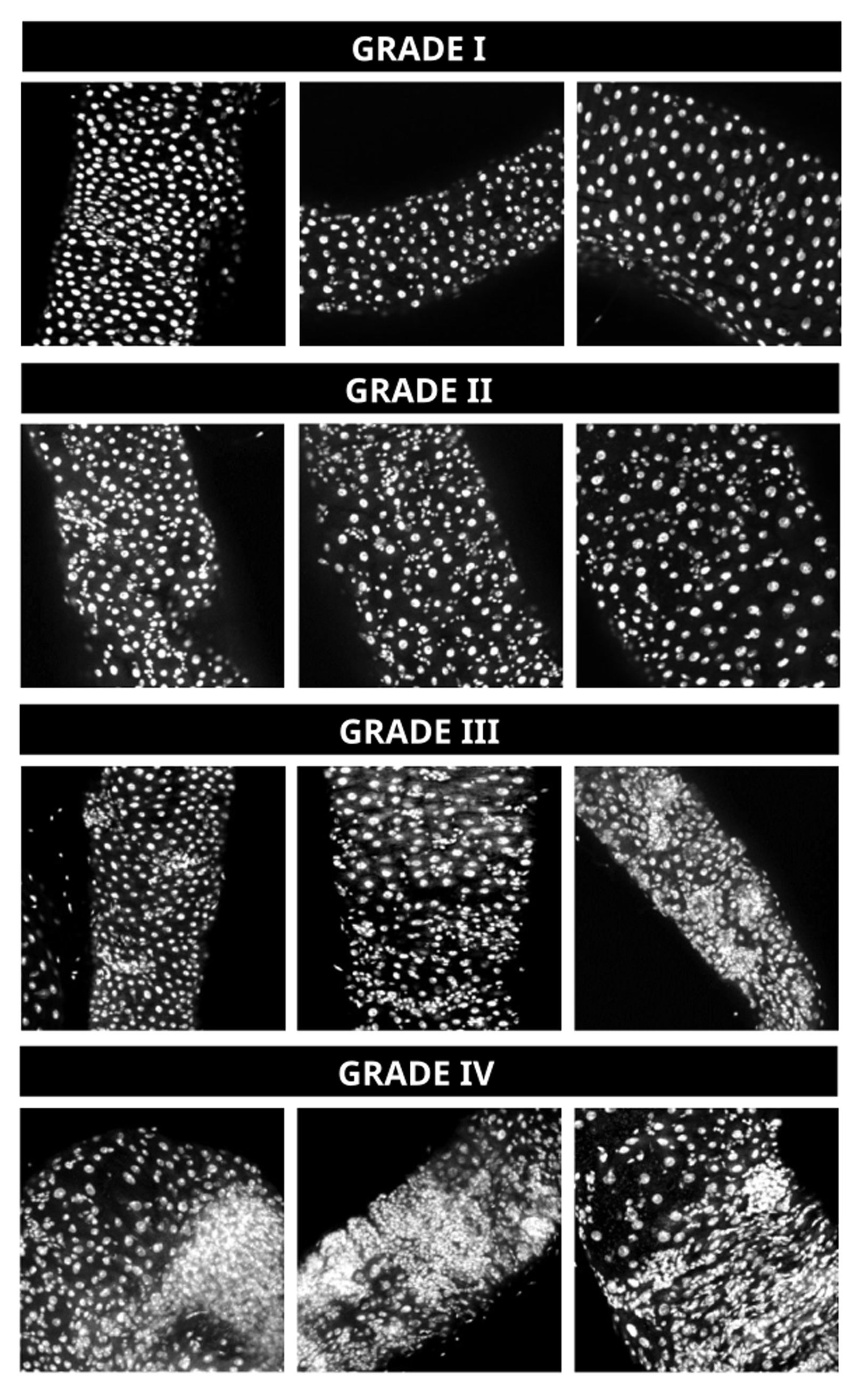
